# Differential effects of AMPA receptor inhibition on seizure-like events in murine and human brain slices

**DOI:** 10.64898/2026.11.10.765159

**Authors:** Franziska Arnold, Alice Podestà, Ran Xu, Julia Onken, Ulrich-Wilhelm Thomale, Angela M. Kaindl, Thomas Sauvigny, Thilo Kalbhenn, Matthias Simon, Christian Bien, Henrik Alle, Jörg R. P. Geiger, Martin Holtkamp, Andrew J. R. Plested, Pawel Fidzinski

## Abstract

Epilepsy is a frequent neurological condition with a worldwide prevalence of 0.5–1%. Current therapies are limited by adverse effects as well as drug resistance. This highlights the need for novel, target-specific antiseizure medications with improved efficacy and fewer side effects.

AMPA (α-amino-3-hydroxy-5-methyl-4-isoxazolepropionic acid) receptors represent a therapeutic target as demonstrated by perampanel, an AMPA receptor antagonist used clinically as antiseizure medication. AMPA receptors typically mediate fast excitatory neurotransmission. Their properties are modulated by auxiliary proteins that can alter receptor activation, deactivation and desensitization, including converting them into slow AMPA receptors.

In this study, we tested different AMPA receptor inhibitors, including perampanel, for their ability to suppress seizure-like events (SLE). SLE were recorded with extracellular electrodes in cortical brain slices from healthy mice and from epilepsy patients who underwent resective surgery. SLE were induced with 4-aminopyridine (4-AP) or a combination of 4-AP and high-potassium. We found that seizure-like events in murine tissue appeared more frequently, with higher amplitude and spike frequency, but were briefer than in human tissue. GYKI53655, NBQX and perampanel were equally effective in inhibiting SLE in murine und human brain slices. In contrast, the antagonist LY3130481, specific for the AMPA receptor auxiliary protein γ-8, showed only weak efficacy in murine tissue, but blocked of SLE in human tissue almost completely.

The species-specific differences we found for LY3130481 highlight the benefit of using human tissue in preclinical research and underline that auxiliary subunits remain a promising therapeutic target for the development of new antiseizure medications.

**Highlights:**

- SLE in murine tissue display higher amplitude and frequency
- SLE in human tissue have longer duration
- The AMPA antagonists GYKI53655, NBQX and perampanel inhibit SLE in both species
- The TARP γ-8 inhibitor LY3130481 has low efficacy to inhibit SLE in murine tissue
- The TARP γ-8 inhibitor LY3130481 nearly abolishes SLE in human tissue

## 1. Introduction

Epilepsy, with a prevalence of 0.5-1%, is one of the most common chronic neurological disorders (1, 2). It is characterized as a predisposition to recurrent, unprovoked seizures, typically resulting from an imbalance between excitation and inhibition (3–6). The primary treatment approach involves anti-seizure medication (ASM) which can achieve seizure freedom in many patients. However, approximately 30% of individuals with epilepsy suffer from drug resistance, defined by the International League Against Epilepsy (ILAE) as failure to achieve seizure freedom by two tolerated ASM at adequate dose used individually or in combination (7, 8). For these patients, epilepsy surgery remains as one of the last therapeutic options.

ASMs are categorized based on their mechanism of action. Established ASMs aim at various targets including voltage-dependent sodium or calcium channels, GABA (γ-aminobutyric acid) receptors, glutamatergic receptors or presynaptic transmitter release (9, 10). Given the significant prevalence of drug-resistant epilepsy, there is need to develop novel ASM with alternative and more specific mechanisms of action. A possible target for new ASM are AMPA (α-amino-3-hydroxy-5-methyl-4-isoxazolepropionic acid) receptors (11). AMPA receptors are glutamate receptors ion channels and typically mediate fast excitatory neurotransmission (12, 13) in the central nervous system. It has been shown that AMPA receptors play a key role in initiation and synchronization of epileptic discharges while NMDA (N-methyl-D-aspartate) receptors prolong synchronized bursts (11). To date, perampanel is the only AMPA receptor antagonist approved as add-on therapy for focal and primary generalized epileptic seizures (14, 15). Although perampanel is effective against seizures, it is associated with relevant adverse effects, including cerebellar symptoms such as ataxia, dysarthria or balance disorders, as well as drowsiness and headaches. These side effects highlight the potential benefits of developing more selective AMPA receptor modulators with better safety profile.

AMPA receptors consist of four subunits (GluA1-4), which can assemble in different combinations. Native AMPA receptors can form protein complexes with different transmembrane auxiliary proteins (3). Among these, transmembrane AMPA Receptor Regulatory Proteins (TARPs) are prominent in the hippocampus, neocortex and striatum (16). The formation of complexes with auxiliary proteins enables functional diversity by changing receptor properties such as activation, deactivation and desensitization (17–21). One AMPA receptor can bind multiple auxiliary proteins simultaneously (22), but whether such complexes have distinct properties is poorly understood. The expression of TARP subtypes differs across brain regions. γ-8, for example, is strongly expressed in the hippocampus but largely absent from the cerebellum (23, 24). In 2016, Kato et al. described LY3130481, an AMPA receptor antagonist that specifically inhibits AMPA receptors that are in complex with TARP γ-8. Since TARP γ-8 is not expressed in the cerebellum, such specificity may help to avoid extrapyramidal motor side effects, which were described for perampanel (25).

In addition to canonical fast AMPA receptor signalling, accumulating evidence suggests the existence of slow AMPA receptors in the brain characterized by slow kinetics and reduced desensitization (26). The high open probability and increased conductance of these variant AMPA receptors (27) result in increased excitability, with consequent potential for greater seizure susceptibility in experimental models (26). To date, slow AMPA receptor mediated currents have been reported in a subset of mouse CA1 principal neurons and to a lesser extent in CA3 principal neurons. The frequency and amplitude of slow AMPA receptor-mediated currents increase when TARP γ-8 is overexpressed (28). In mouse hippocampus, another auxiliary protein, CNIH3, is decisive for slow AMPA receptor expression (29). In this study, we aimed to compare the efficacy of AMPA receptor antagonists with distinct specificity regarding their ability to inhibit epileptiform activity *in vitro*. We used both murine and human brain tissue to investigate species-specific differences.

## 2. Methods

### 2.1 Slice preparation – mice

All animal procedures complied with the German Animal Welfare Act and the European Directive 2010/63/EU on protection of animals used for scientific purposes. All experiments were approved by the Institutional Animal Welfare Officer and the local regulatory authority (Landesamt für Gesundheit und Soziales, Berlin, Germany; license number T-Ch 001021). Acute hippocampal-entorhinal cortex slices were obtained from male C57BL/6 mice aged 3-4 weeks. The preparation was carried out as described previously (30): Mice were anaesthetized with isoflurane (4% in 100% O2) and decapitated. The brain was rapidly removed (<60s) and placed in ice-cold, sucrose-based artificial cerebrospinal fluid (aCSF) saturated with carbogen (Lindner Gase) and containing (in mM): NaCl 87, NaHCO_3_ 25, NaH_2_PO_4_ 1.25, KCl 2.5, CaCl_2_ 0.5, MgCl_2_ 7, glucose 10 and sucrose 75. After removal of the cerebellum and hemispheric separation, the hemispheres were cut horizontally in 450 µm thick slices using a vibratome (VT1200S, Leica). Slices included temporal (TC), perirhinal (PC) and entorhinal (EC) cortex, subiculum (SUB), dentate gyrus (DG) and cornu ammonis (CA) regions. The sucrose-based aCSF was used during slicing under continuous carbogenation. Slices were then transferred and stored in an interface chamber with prewarmed (35°C), carbogenated aCSF containing (in mM): NaCl 129, NaH_2_PO_4_ 1.25, CaCl_2_ 1.6, KCl 3, MgSO_4_ 1.8, NaHCO_3_ 21, glucose 10) and allowed to recover for at least 30 minutes before electrophysiology.

### 2.2 Slice preparation – human brain tissue

Human brain tissue was obtained from epilepsy patients that underwent resective surgery. All patients gave informed written consent prior to surgery. Tissue samples originated from three neurosurgical departments: Charité-Universitätsmedizin Berlin; University Medical Center Hamburg Eppendorf and Bethel Klinikum Bielefeld. The use of resected tissue for research purposes was approved by local ethics committees (Ethical vote numbers: Berlin: EA1_111_14 and EA2/064/22; Bielefeld: 2020-517-f-S; Hamburg: 2023-200674-BO-bet). Tissue was taken from 24 patients who were between 4 and 59 years old at time of resection (mean 33,5 years) and 37,5% were female (Table 1). The preparation was carried out as previously described (31): Immediately after resection, the tissue was immersed in ice-cold, carbogenated sucrose based aCSF (containing in mM: NaCl 87, NaHCO_3_ 25, NaH_2_PO_4_ 1.25, KCl 2.5, CaCl_2_ 0.5, MgCl_2_ 3, glucose 10, sucrose 75) and transported to the laboratory in Berlin. Before slicing, the meninges were carefully removed with blunt forceps, the tissue was trimmed, glued to a vibratome chamber and cut into 450 µm thick horizontal slices in carbogenated, sucrose based aCSF. After slicing, tissue underwent recovery in sucrose-based aCSF at 35°C in a custom-made interface system for at least 30 minutes. After recovery, slices were stored at room temperature. All procedures were conducted under near-sterile conditions. If tissue was resected at external sites (Hamburg or Bielefeld), slices were prepared remotely, put in custom transport containers filled with carbogenated sucrose-aCSF and transferred to Berlin, where they were transferred again to the same type of interface system used for recovery.

**Table 1.**

| **No.** | **Age** | **Sex** | **Brain area of resection** | **Diagnosis** | **Last ASM** |
|---|---|---|---|---|---|
| 1 | 4 | male | Temporal pole resection | FCD IIIa | Lamotrigine, clobazam |
| 2 | 31 | male | Temporal pole resection | Encephalocele | Levetiracetam, lamotrigine |
| 3 | 58 | male | Temporal pole resection | Hippocampal sclerosis | Brivaracetam, perampanel, zonisamide |
| 4 | 57 | male | Temporal pole resection | Encephalocele | Gabapentin, lamotrigine |
| 5 | 24 | male | Temporal pole resection | Cavernoma (cavernous hemangioma) | Lacosamide |
| 6 | 58 | male | Temporal pole resection | Hippocampal sclerosis | Lamotrigine |
| 7 | 38 | female | Temporal (extended lesionectomy ) | Meningoencephalocele with gliosis | Brivaracetam, lacosamide |
| 8 | 37 | female | Temporal pole resection | Hippocampal sclerosis | Lamotrigine |
| 9 | 48 | female | Temporal pole resection | Hippocampal sclerosis left, temporal encephalocele | Lamotrigine |
| 10 | 27 | male | Temporal pole resection | Hippocampal encephalitis | Lacosamide, brivaracetam |
| 11 | 15 | female | Temporal (extended lesionectomy) | Ganglioglioma WHO I | Brivaracetam, lamotrigine |
| 12 | 16 | female | Occipital (lesionectomy) | T2 hyperintensitiy occipital right - FCD suspected | Clobazam, perampanel, cenobamate |
| 13 | 27 | female | Temporal pole resection | Meningocele | Lamotrigine, brivaracetam, lacosamide |
| 14 | 58 | male | Temporal pole resection | Hippocampal sclerosis | Lacosamide, perampanel |
| 15 | 49 | female | Temporal pole resection | Suspected gangliolioma, apart from it gliosis | Levetiracetam, lamotrigine, perampanel |
| 16 | 36 | female | Temporal pole resection | Hippocampal sclerosis | Lacosamide |
| 17 | 22 | male | Temporal pole resection | Gangliogliom | Clobazam, perampanel, cenobamate |
| 18 | 35 | male | Temporal pole resection | Arachnoidal cyst | Lamotrigine |
| 19 | 13 | male | Temporal pole resection | Central neurocytoma | Brivaracetam, lamotrigine |
| 20 | 37 | male | Temporal pole resection | Hippocampal sclerosis | Oxcarbazepine, brivaracetam, lamotrigine |
| 21 | 21 | female | Temporal pole resection | Hippocampal sclerosis | Levetiracetam, lacosamide, clobazam |
| 22 | 32 | male | Temporal pole resection | Ganglioglioma WHO I | Lacosamide, perampanel |
| 23 | 32 | female | Temporal pole resection | Hippocampal sclerosis | Eslicarbazepin, perampanel |
| 24 | 28 | male | Temporal pole resection | Hippocampal sclerosis | Levetiracetam, lacosamide |

### 2.3 Electrophysiological recordings

Extracellular local field potentials (LFP) were recorded using glass electrodes (1-2 MΩ) filled with 154 mM NaCl. Electrodes were positioned in layer II/III of the entorhinal cortex in mice or the medial temporal gyrus of the temporal neocortex (NC) in human slices. Field potential recordings were performed using custom-built amplifiers (10x), a Micro 1401 mk II AD converter (Cambridge Electronic Design Limited, Cambridge, UK) and Spike2 software (v7.00). To induce seizure like events (SLE) in mouse slices, 100 µM 4-aminopyridine (4-AP) was applied to inhibit potassium channels. Human slices were exposed to 10 mM K^+^-aCSF (in mM: NaCl 122, NaH_2_PO_4_ 1.25, CaCl_2_ 1.6, KCl 10, MgSO_4_ 1.8, NaHCO_3_ 21, glucose 10) in combination with 200 µM 4-AP. 4-AP is a non-selective potassium channel blocker. aCSF with elevated potassium levels leads to cell depolarization, prolongs action potential duration and can induce bursting activity in neurons (32). For one set of experiments in human tissue SLEs were induced by 0 mM Mg^2+-^aCSF (in mM: NaCl 130.8, NaH_2_PO_4_ 1.25, CaCl_2_ 1.6, KCl 3, NaHCO_3_ 21, glucose 10) in combination with 10 μM gabazine (GBZ) and 10 μM CGP 55845 to activate NMDA receptors and block GABA-A and GABA-B-receptors (32). SLE were defined as field potential deflections of ≥ 0.5 mV, ≥ 10 s duration and the presence of superimposed ripple-like discharges. After the occurrence of the first SLE, a 30 min baseline was recorded. If no SLE occurred within 45 minutes of induction, the slice was considered as not responsive and excluded. After baseline recording, the investigated substances were applied and allowed to equilibrate for 15 minutes (40 min for LY3130481 due to slow tissue penetration (25)). The treatment period was recorded for 30 minutes, followed by washout and another 30 min recording period. The following drugs were used: LY3130418 inhibits AMPA receptors associated with TARP γ-8 and thus might also inhibit slow AMPA currents (25). NBQX is a competitive glutamate receptor antagonist targeting kainate receptors and canonical fast AMPA currents (28). GYKI53655 is a non-competitive antagonist that inhibits both classical and slow AMPA currents ((26), (33)). Perampanel is a non-competitive, AMPA receptor antagonist (34, 35) that also has some inhibitory activity against kainate receptors (36).

### 2.4 Data processing and analysis

SLE were analysed using custom-written scripts in Matlab (v8.6, R2015b, MathWorks, Natick, MA, USA). Analysed SLE parameters included incidence, duration, amplitude, population spike frequency and population spike count. Statistical analysis was performed with GraphPad Prism (v10.3.1, GraphPad Software, San Diego California USA). Data were tested for normality with D’Agostino & Pearson omnibus test. Group comparisons were made using the Kruskal-Wallis test followed by Dunn’s multiple comparisons test or Mann-Whitney-Test when only one treatment group was compared to control. In the 0 mM Mg^2+^-aCSF +CGP 10 µM + 10 µM GBZ condition the intervention and washout period was compared to baseline therefore the Friedman test was used followed by Dunn’s multiple comparison test. P-values <0.05 were considered statistically significant. Figures were created with Inkscape (v1.2.1, The Inkscape Project). Pictograms (mouse and human) were taken from Flaticon.com.

## 3. Results

### 3.1 SLE induction in mouse and human brain tissue *in vitro*

To assess the effects of AMPA receptor antagonists on ictogenesis, established models for SLE induction *in vitro* were used (Figure 1). In murine tissue, 4-AP model reliably induced SLE activity (Figure 1C, E). In contrast, SLE induction in human brain tissue was more challenging. To facilitate comparability to the murine model, we used a model which combines 4-AP with elevated extracellular potassium (10 mM). Using this method yielded reproducible SLE in human slices, albeit with a lower SLE incidence compared to murine tissue (Figure 1E). SLE in human brain tissue had lower amplitudes, longer durations and lower spike frequencies (Figure 1E). Under control conditions, SLE activity in human slices was stable for approximately two hours, precluding meaningful analysis of longer-lasting washout effects, which therefore are not shown in subsequent figures.

**Figure 1.**
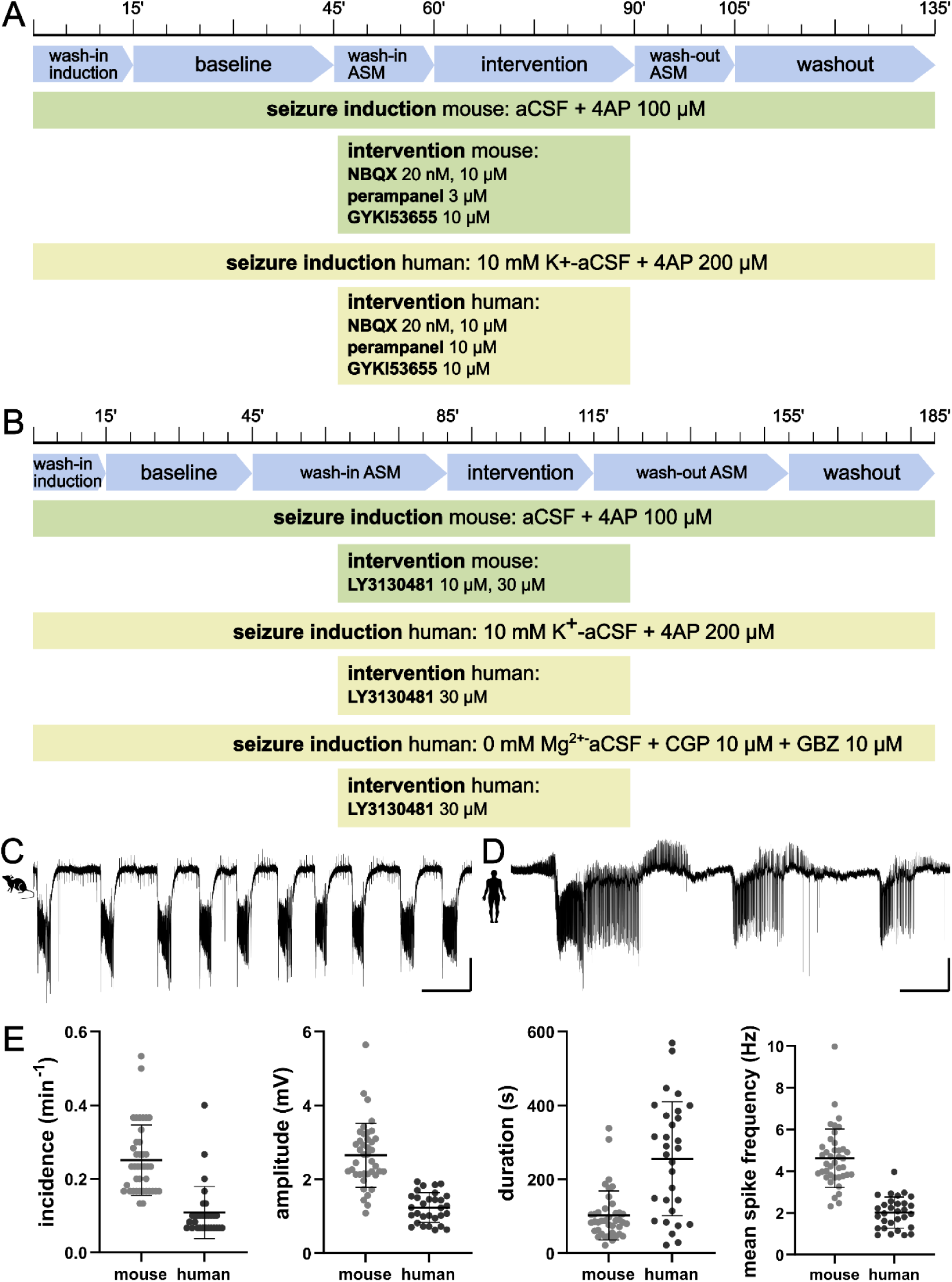
Flow diagram of experimental protocol in mice and human brain tissue and SLE induction in mouse and human brain tissue in vitro. (A) Protocol for experiments with NBQX, perampanel and GYKI53655. Each experiment consists of three recording periods à 30 minutes with wash-in and wash-out phases of 15 minutes in between. (B) Protocol for experiments with LY3130481 and controls. Experiments have the same recording periods but the wash-in and wash-out of the ASM was extended to 40 minutes. (A-B) In the mouse model SLEs were induced by adding 4-AP to aCSF, in human tissue 10 mM K^+^-aCSF was combined with 4-AP or 0 mM Mg^2+-^aCSF was combined with CGP and Gabazine. (C-D) Representative LFP recordings in mouse entorhinal cortex (C) and human neo cortex layer II -III using the 10mM K^+^-aCSF + 4AP induction method (D). Traces show baseline period, scale bar: 200 s, 1 mV. (E) Comparison of SLE incidence, amplitude, duration and mean spike frequency (from left to right) during baseline period between mice and human brain tissue using the 10 mM K^+^-aCSF + 4AP induction method. Scatterplots show mean values ± standard deviation. Sample sizes: mouse: 40 slices, human: 30 slices.

### 3.2 NBQX, GYKI53655 and perampanel abolish SLE in mouse and in human brain tissue

To explore the role of canonical fast AMPA receptors, we used NBQX, a competitive antagonist of AMPA and kainate receptors with preferential effects on fast AMPA receptor mediated currents. In murine slices, 10 µM NBQX fully blocked SLE during application (p<0,01) (Figure 2A) and during washout (p<0,01) (Figure S1). Due to complete block, other SLE parameters could not be analysed in NBQX. We next tested additional AMPA receptor antagonists. GYKI53655 (10 µM), a selective, non-competitive AMPA receptor inhibitor which does not act on kainate receptors, completely blocked murine SLE during application (p<0,01). Perampanel (3 µM), a selective, non-competitive AMPA receptor antagonist, which again also affects kainate receptors, almost completely blocked SLE (normalized SLE incidence: 0.03±0,06; p<0,01) (Figure 2A). For both GYKI53655 and perampanel, SLE activity recovered during washout (Figure S1). In human tissue, 10 µM NBQX strongly but not completely inhibited SLE (normalized SLE incidence 0.01±0.03; p<0,01) while GYKI53655 (p<0.01) and perampanel at 10 µM as per literature completely blocked SLE (37) (p<0.01) (Figure 2B).

**Figure 2.**
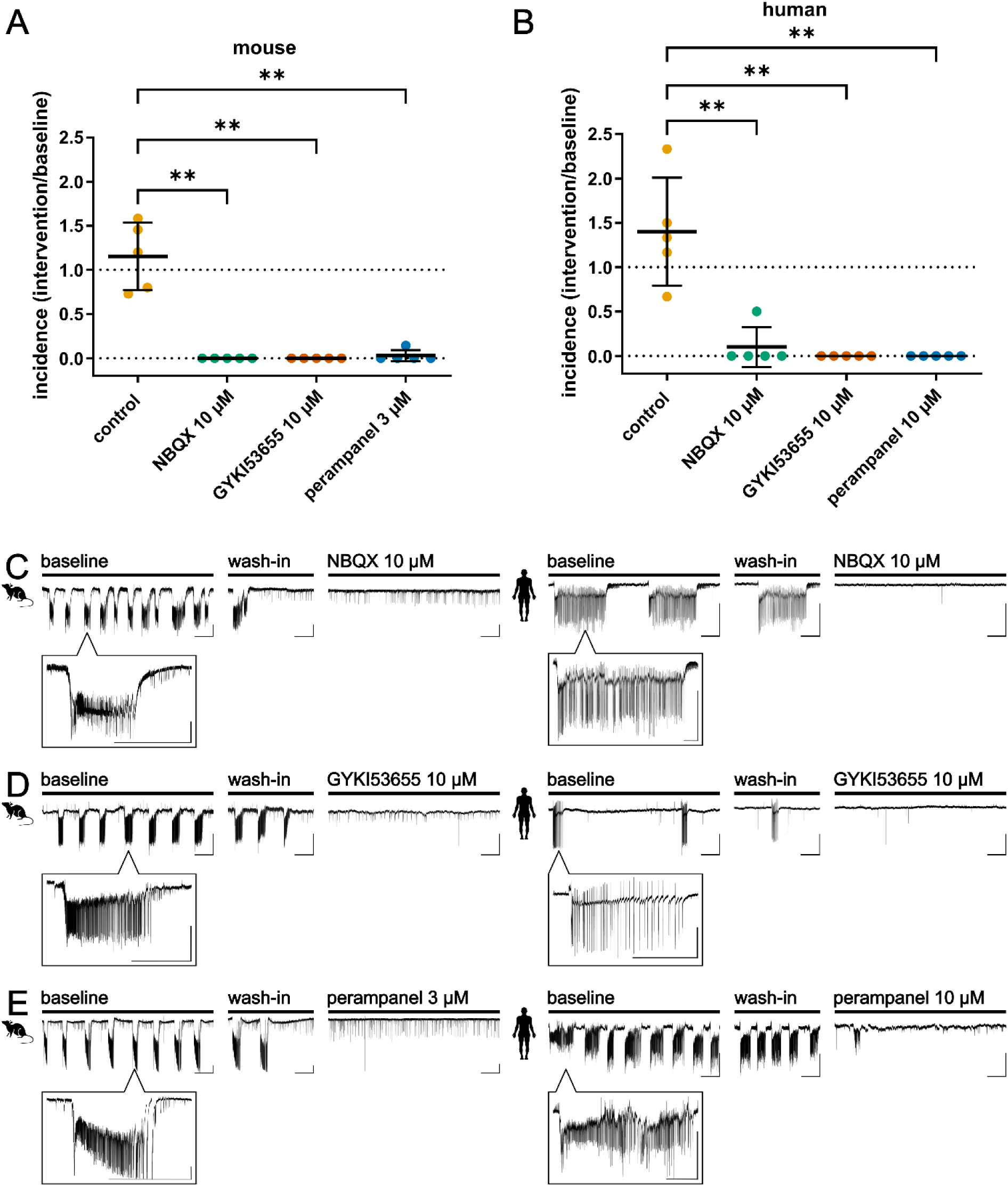
NBQX, GYKI53655 and perampanel block SLEs in mouse and in human brain tissue. (A-B) Scatterplots of SLE-incidence during intervention period normalized to baseline (intervention/baseline) for NBQX, GYKI53655 and perampanel in (A) mouse and (B) human tissue. In human tissue the using the 10 mM K^+^-aCSF + 4AP induction method was used. Data are shown as mean ± standard deviation. Asterisks mark statistically significant differences assessed by Kruskal-Wallis test and Dunn’s multiple comparisons test (*p ≤ 0.05, **p ≤ 0.01, ***p ≤ 0,001). (C-E) Representative traces of LFP-recordings for (C) NBQX, (D) GYKI53655 and (E) perampanel. Recordings in mice are shown on the left, recordings in human brain tissue on the right, scale bar: 200 s, 1 mV. Enlargements of one individual SLE for each recording period, scale bar: 60 s, 1 mV.

### 3.3 Low concentrations of NBQX do not reduce of SLE incidence in murine or human slices

At very low concentrations (below 100 nM), NBQX has been shown to inhibit classical fast AMPA currents, whilst also reducing desensitization and sparing or even potentiating slow AMPA receptor responses (28, 38). To investigate whether differential modulation of fast and slow AMPA receptor affects ictogenesis, we examined the effects of NBQX at 20 nM on SLE. NBQX (20 nM) did not significantly alter SLE incidence in either mouse brain slices (normalized SLE incidence 0.68±0.33, p>0.9) or human brain slices (normalized SLE incidence 0.98±0.32, p=0.67) when compared to control (Figure 3A). In mouse slices, SLE amplitude, duration and mean spike frequency remained likewise unaltered (Figure S2A). In human brain slices, SLE duration was reduced in the control group relative to baseline whereas it remained comparable to baseline level in the NBQX group. In other words, SLE duration was higher (p=0.0078) in the NBQX group than in the control group (normalized SLE duration 0.86±0.26 for NBQX vs. 0.35±0,04 in the control group) (Figure S2B). SLE amplitude and mean spike frequency did not change. Thus, in human slices, 20 nM NBQX did not prevent SLE initiation but was associated with relatively prolonged SLE duration, raising the possibility that slow AMPAR-mediated currents contribute to SLE maintenance.

**Figure 3.**
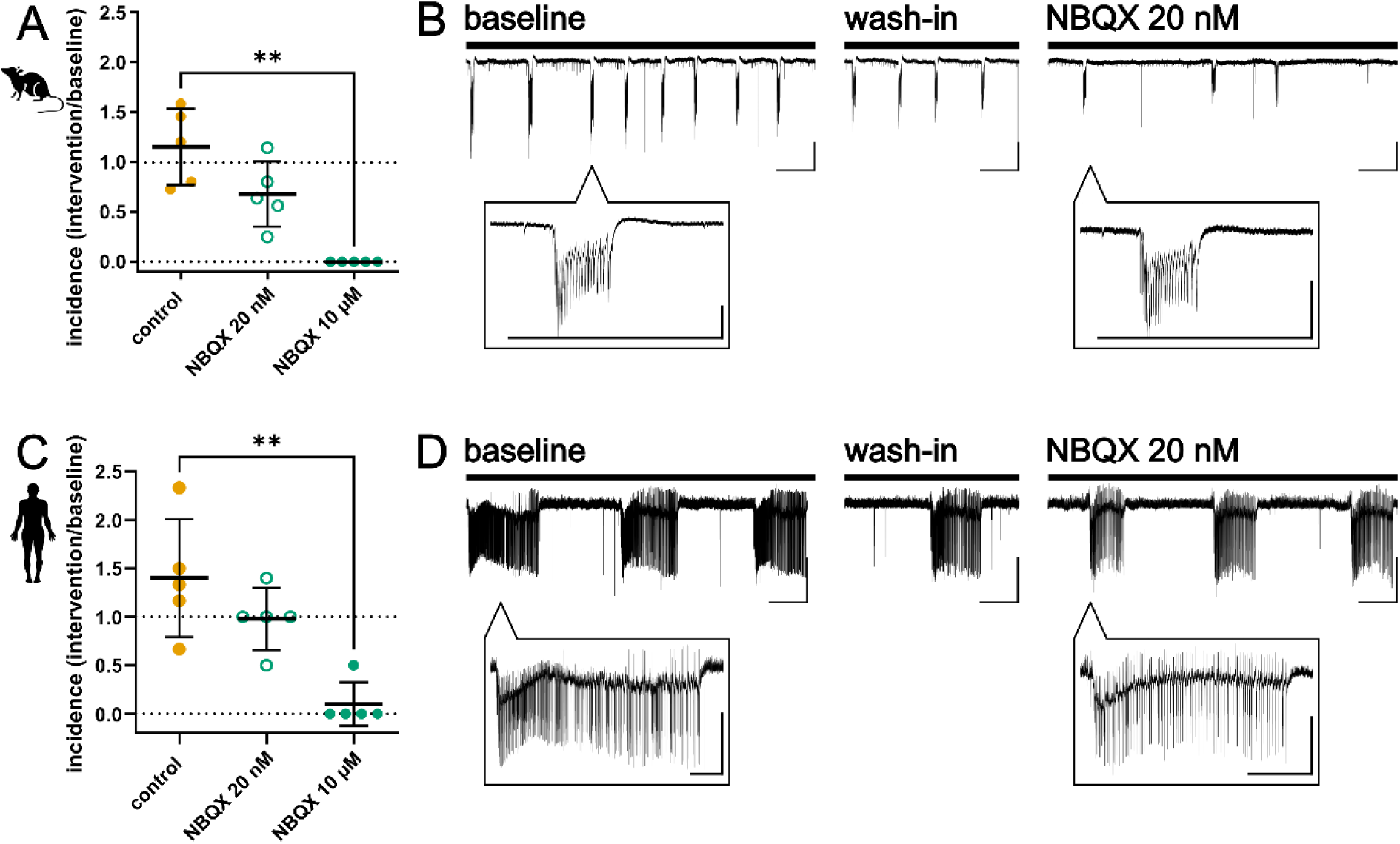
Very low concentrations of NBQX show no significant reduction of SLE incidence in mouse and human brain slices. SLEs in human tissue were induced by 10 mM K^+^-aCSF + 4AP. (A, C) Scatterplots of SLE-incidence during intervention period normalized to baseline (intervention/baseline) in (A) mouse model and (B) human tissue compared between control group, NBQX 20 nM and NBQX 10 µM. Data are shown as mean ± standard deviation. Asterisks mark statistically significant differences assessed by Kruskal-Wallis test and Dunn’s multiple comparisons test (*p ≤ 0.05, **p ≤ 0.01, ***p ≤ 0,001). (B, D) Representative traces of LFP-recordings for NBQX 20 nM in (B) mouse and (D) human brain tissue, scale bar: 200 s, 1 mV. Enlargements of one individual SLE for each recording period, scale bar: 60 s, 1 mV.

### 3.4 LY3130481 fully inhibits SLE in human tissue but only partially in mouse tissue

To assess the involvement of TARP γ8 in neocortical ictogenesis *in vitro*, we investigated the effects of the γ8-specific AMPA receptor inhibitor LY3130481 on mice and human SLE. As expected (25), brief application of LY3130481 (10 µM) for 15 min in mouse slices did not alter SLE activity. Extending the incubation to 40 min resulted in a reduction of SLE incidence, but only in a subset (30%) of murine slices. The responder rate – defined as ≥ 50% reduction of SLE incidence – reached 40% at the concentration of 30 µM (Figure 4C). 30 µM LY3130481 reduced normalized SLE incidence to 0,48±0,19 (p<0.05) while in the control group normalized SLE incidence remained unchanged (1.15±0.38). Overall, using 10 µM LY3130481 failed to significantly reduce SLE incidence (0.90±0.57; p=0.6) (Figure 4A). Other SLE parameters (amplitude, duration, spike frequency and spike count) remained likewise unchanged (Figure S1). Since 30 µM proved to be effective in mice, we chose this concentration for experiments in human tissue. In contrast to the partial effect in murine tissue, 30 µM LY3130481 gave a strong and almost complete block of SLE in human tissue with a normalized SLE incidence of 0.10±0.22 (p < 0.01) (Figure 4B, C, F). As previously, the effect of LY3130481 to abolish SLE in human tissue was too strong to measure SLE characteristics in more detail.

**Figure 4.**
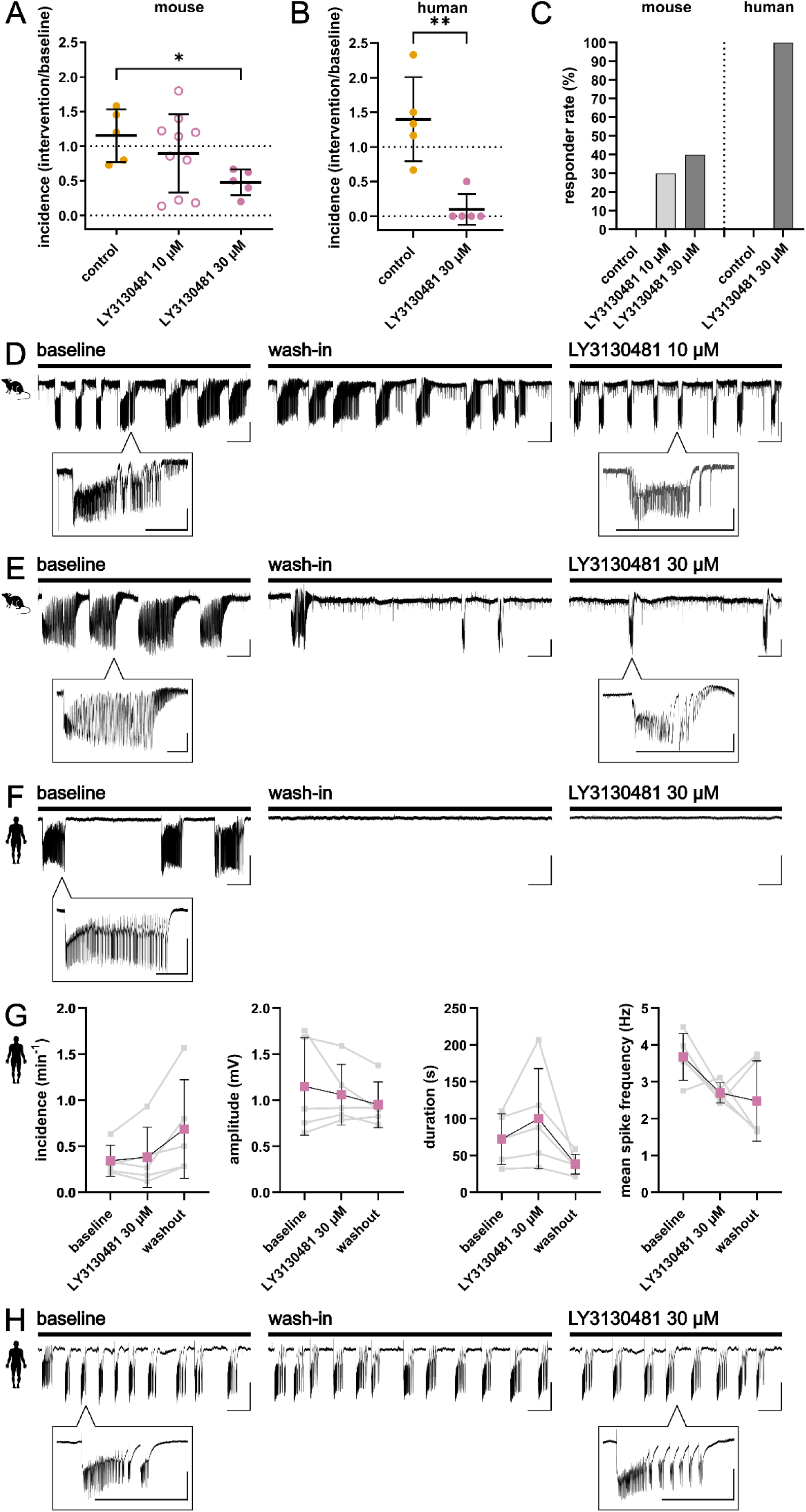
LY3130481 completely blocks SLEs in human tissue in the 10 mM K^+^-aCSF + 4AP model, doesn’t block SLEs in the 0 mM Mg^2+^-aCSF + CGP + Gabazine model and shows only partial effect in mouse 4-AP model. (A) Scatterplot of SLE-incidence during intervention period normalized to baseline (intervention/baseline) in mouse model compared between control group, LY3130481 10 µM and LY3130481 30 µM. Data are shown as mean ± standard deviation. Asterisks mark statistically significant differences assessed by Kruskal-Wallis test and Dunn’s multiple comparisons test (*p ≤ 0.05). (B) Scatterplot of SLE-incidence during intervention period normalized to baseline (intervention/baseline) in human tissue compared between control group and LY3130481 30 µM. SLEs were induced using the 10 mM K^+^-aCSF + 4AP induction method. Data are shown as mean ± standard deviation. Asterisks mark statistically significant differences assessed by Mann-Whitney test (*p ≤ 0.05). (C) Bar chart of responder rate defined as a reduction of SLE-incidence during intervention period of at least 50% compared to baseline. (D-F) Representative traces of LFP-recordings for each condition: (D) mouse LY3130481 10 µM, (E) mouse LY3130481 30 µM, (F) human LY3130481 30 µM, scale bar: 200 s, 1 mV. Enlargements of one individual SLE for each recording period, scale bar: 60 s, 1 mV. (G) Effects of LY3130481 30 µM on SLE incidence, amplitude, duration and mean spike frequency (from left to right) *in vitro* in human brain tissue induced by 0 mM Mg^2+^-aCSF + CGP 10 µM + GBZ 10 µM. Scatterplots represent individual recordings, black line represents mean ± standard deviation. Analysis was done by using Friedman test and Dunn’s multiple comparisons test. (H) Representative trace of one LFP recording using the 0 mM Mg^2+^-aCSF + CGP + Gabazine model and LY3130481 30 µM. Scale bar: 200 s, 1 mV. Enlargements of one individual SLE for each recording period, scale bar: 60 s, 1 mV.

Using 0 mM Mg^2+^-aCSF in combination with 10 μM gabazine (GBZ) and 10 μM CGP 55845 induces SLE by allowing NMDA receptor currents and blocking inhibition from GABA-A and GABA-B-receptors. In contrast to the high potassium aCSF / 4-AP model, SLEs induced by 0 mM Mg^2+-^aCSF were not affected by LY3130481, suggesting that in this model SLE maintenance is not modulated by AMPA receptors containing the TARP γ8 subunit. (Figure 4H).

## 4. Discussion

In the present study, we investigated the effects of a panel of AMPA receptor antagonists on epileptiform activity, represented by seizure-like events in extracellular recordings of both mouse and human brain slices. We focused on LY3130481, a selective modulator of AMPA receptors in complex with TARP γ8, with the aim of exploring its potential of subtype-specific AMPA receptor inhibition. Our key findings are: 1) SLEs in murine tissue occurred at higher incidence and with greater amplitude and spike frequency than SLEs in human tissue, while the latter were longer in duration. 2) LY3130481 efficacy to inhibit SLE in murine tissue was low, with only 40% of responsive slices at the highest tested concentration, while 3) the same dose of LY3130481 nearly abolished SLE with a 100% responder rate in human slices. 4) Other AMPA antagonists, including GYKI53655, NBQX and perampanel strongly inhibited SLE in both species.

### 4.1 LY3130481

LY3130481 (also known as CRC-611 and, more recently, ES-481) was first described in 2016 as a forebrain-specific drug candidate with antiepileptic action (39). LY3130481 binds between transmembrane helices three and four on TARP γ-8 but does not dissociate the auxiliary protein from the AMPA receptor (40). It is not competitive with glutamate, and it does not affect NMDA, kainate or metabotropic glutamate receptors (25). LY3130481 was effective in reducing seizure (or seizure-like) activity in several models, including single cell experiments, mouse and rat *in vivo* models and AMPA receptors from an epilepsy patient (25). Another study used whole cell recordings of the GluA2 subunit containing TARP γ-8 expressed in HEK293T cells and was able to show that LY3130481 suppressed resensitization and reduced the equilibrium current but did not reduce the amplitude of the peak current (40). In our study we tested the effect of LY3130481 on SLEs induced *in vitro* mouse tissue and tissue samples from five epileptic patients. Our main finding was that LY3130481 was less effective in mice than in human tissue. This species-specific efficacy could be due to a lower expression of TARP γ-8 in certain murine neurons, although previous data suggests similar TARP γ-8 expression levels in mice, rats and humans with the protein being preferentially expressed in the forebrain and enriched in the hippocampus (41). In mice, TARP γ-8 expression does not change much over the lifespan (42) whereas in rats expression increases with age (43). It is not known whether TARP γ-8 expression in the human brain changes with age or due to epilepsy. Witkin et al. reported lower radioligand binding density in human brain compared to rodent brain (44), although this finding seems at odds with our functional data showing higher sensitivity to LY3130481 in human tissue. Binding density alone may not directly reflect functional inhibition and may vary between cortical regions and pathological states. It is also possible that human epileptic tissue exhibits different expression of slow AMPA receptor complexes, particularly relevant to the putative influence of TARP γ-8. Prior studies have shown that overexpression of TARP γ-8 increases the frequency and amplitude of slow AMPA currents in rodent hippocampus (28). Such slow kinetics in pyramidal cells seem likely to lead to increased excitability and seizure susceptibility (26). The nearly-complete inhibition of SLEs by LY3130481 in human slices could be due to pathological enrichment of TARP γ-8-associated AMPA signalling in pharmacoresistant epilepsy.

### 4.2 Comparison with canonical AMPA antagonists

In contrast to the selective effect of LY3130481, other, non-competitive AMPA receptor antagonists such as GYKI53655 and perampanel, as well as the competitive antagonist NBQX, inhibited SLE in both species. GYKI53655, a non-competitive inhibitor that largely spares kainate receptors, and perampanel, which acts on AMPA but also blocks kainate receptors (36), were highly effective, consistent with previous *in vivo* and clinical findings ((14), (37), (33), (45)). The concentration of perampanel we used in this study is approximately ten times higher than the one usually found in patients treated with the substance (46), which seems to explain the strong effect in human tissue of pharmacoresistant patients. NBQX showed full inhibition in mice and near-complete inhibition in humans, reinforcing its established role in blocking fast AMPA currents. Importantly, NBQX at a very low concentration (20 nM), which is unlikely to affect slow AMPA receptors, but should still inhibit classical AMPAR activity, failed to alter SLE activity, supporting the notion that fast and slow AMPA signalling may be pharmacologically separable. Interestingly, although this concentration NBQX did not affect SLE, the relative prolongation of SLE duration in human slices may indicate that slow AMPAR-mediated currents contribute to the maintenance of established epileptiform activity.

### 4.3 Translational considerations

The differential drug responses between murine and human tissue point to an important translational aspect. While the 4-AP model in young, healthy mice provides a stable and reproducible platform for testing antiseizure agents, it may not adequately reflect the structural and functional alterations found in human tissue. Differences in network architecture, receptor composition, and auxiliary protein expression all contribute to species-specific pharmacodynamics. Moreover, the nearly complete blockade of SLE by LY3130481 in human slices suggests a potential therapeutic advantage over non-selective AMPA receptor antagonists like perampanel, which are associated with cerebellar side effects due to widespread AMPA receptor inhibition. Since TARP γ-8 is largely absent from the cerebellum (23), LY3130481 and other drugs targeting diverse AMPA receptor subtypes may offer region-selective inhibition with improved tolerability.

### 4.4 Limitations

This study has several limitations. First, while the strong effect of LY3130481 in human tissue suggests high expression of TARP γ-8 in the human brain, we did not biochemically quantify TARP γ-8 expression in murine or human brain tissue via western blot or immunostaining. Second, due to limited availability of human specimen, we did not evaluate the full concentration-response range for LY3130481. Testing lower doses in human slices might reveal a therapeutic window or dose-dependency more reflective of clinical conditions. Third, the *in vitro* models applied in this study rather reflect acute-symptomatic seizures than epilepsy since SLE do not occur spontaneously but are triggered pharmacologically. Finally, our data does not reveal information about potential mechanisms as we only used extracellular recordings - intracellular recordings would be required to directly assess the effects of slow AMPA receptor kinetics.

## 4.5 Conclusions

Our findings demonstrate that the TARP γ-8–selective AMPA receptor antagonist LY3130481 suppresses epileptiform activity in human neocortical tissue but has only limited efficacy in mouse tissue. This species-specific difference highlights the importance of using human tissue in preclinical research. The contribution of TARP γ-8–associated AMPA signalling in human ictogenesis suggests that more selective modulators of AMPA receptor subtypes may offer therapeutic benefit with improved safety profiles. Further studies will be needed to address the validation of pathological relevance of AMPA receptor subunits in human seizures and epilepsy and evaluation of other modulators targeting particular auxiliary subunits such as JNJ-61432059.

## Acknowledgments

We would like to thank Mandy Marbler-Pötter and Andrea Wilke for their excellent technical assistance.

## Data Availability

Data are available upon request to the corresponding authors.

## CRediT authorship contribution statement

**Franziska Arnold:** Investigation, Data Curation, Formal Analysis, Visualization, Writing – Original Draft. **Alice Podestà:** Investigation. **Ran Xu:** Resources. **Julia Onken:** Resources. **Ulrich-Wilhelm Thomale:** Resources. **Angela M. Kaindl:** Resources. **Thomas Sauvigny:** Resources. **Thilo Kalbhenn:** Resources. **Matthias Simon:** Resources. **Christian Bien:** Resources. **Henrik Alle:** Project Administration. **Jörg R. P. Geiger:** Project Administration, Funding Acquisition. **Martin Holtkamp:** Conceptualization, Funding Acquisition, Project Administration. **Andrew J. R. Plested:** Conceptualization. **Pawel Fidzinski:** Conceptualization, Methodology, Project Administration, Supervision. All authors critically reviewed and revised the manuscript.

---

**Figure S1.**
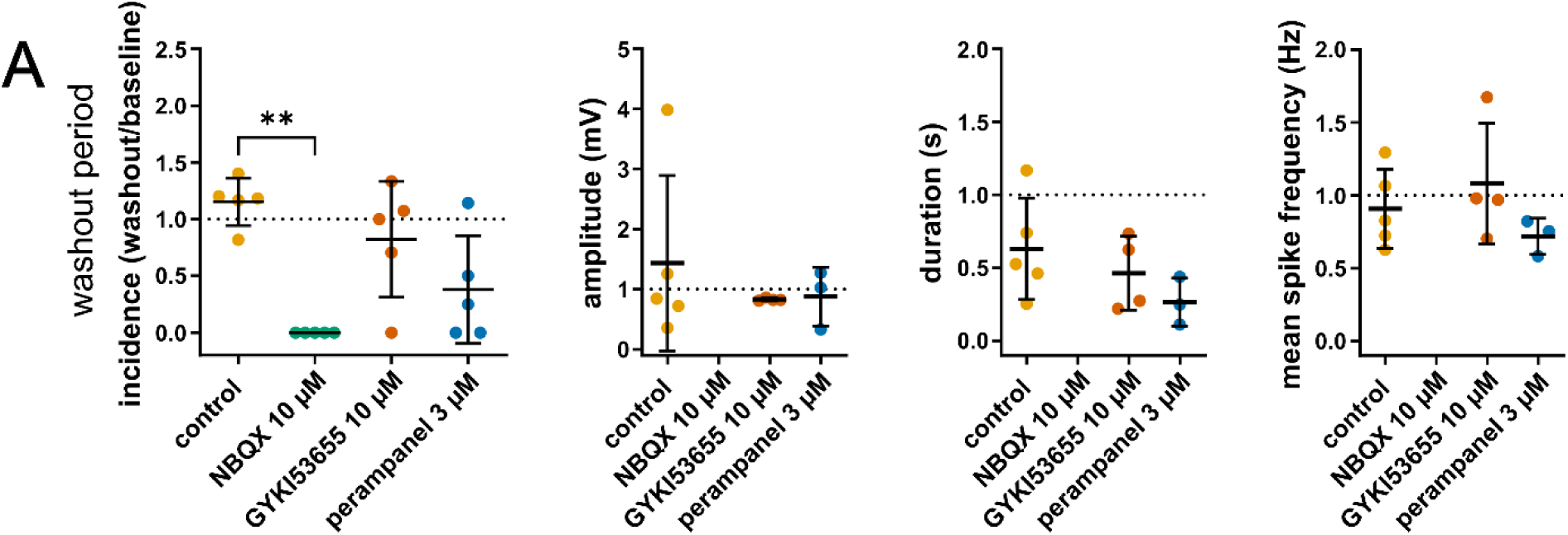
Effects of different AMPA receptor antagonists on seizure-like events *in vitro* in mice induced by 4-AP. Scatterplots show mean values ± standard deviation. All values are normalized to baseline values (washout/baseline). Asterisks mark statistically significant differences assessed by Kruskal-Wallis test and Dunn’s multiple comparisons test (*p ≤ 0.05, **p ≤ 0.01, ***p ≤ 0,001). (A) Scatterplots (from left to right) of SLE incidence, amplitude, duration, mean spike frequency and spike number during washout period compared between control group, NBQX 10 µM, GYKI53655 10 µM and perampanel 3 µM.

**Figure S2.**
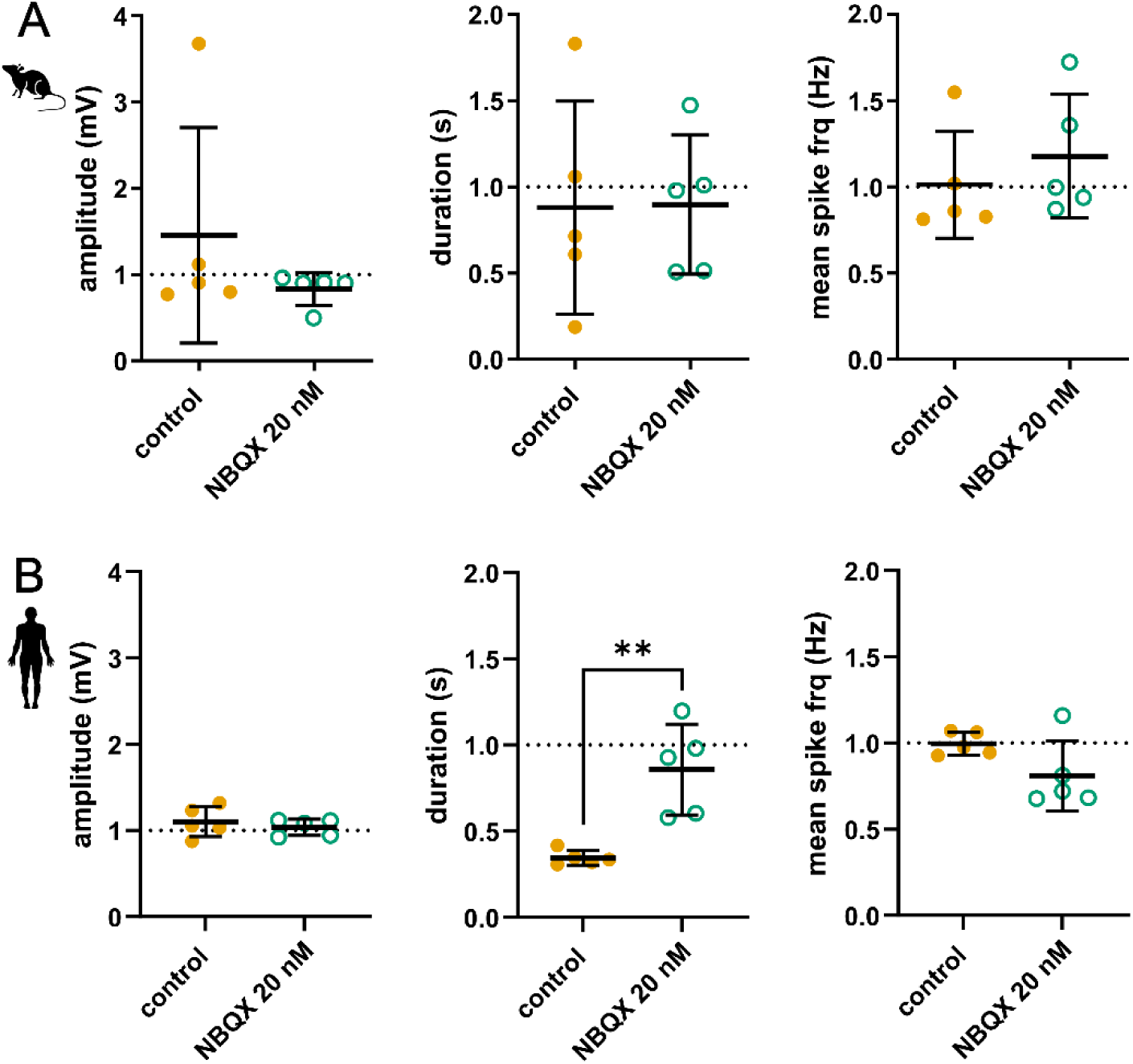
Effects of very low NBQX concentration on SLE amplitude, duration and mean spike frequency (from left to right) in (A) mouse and (B) human brain tissue. Scatterplots show mean values ± standard deviation. All values are normalized to baseline values. Asterisks mark statistically significant differences assessed by Mann-Whitney test (*p ≤ 0.05, **p ≤ 0.01, ***p ≤ 0,001).

**Figure S3.**
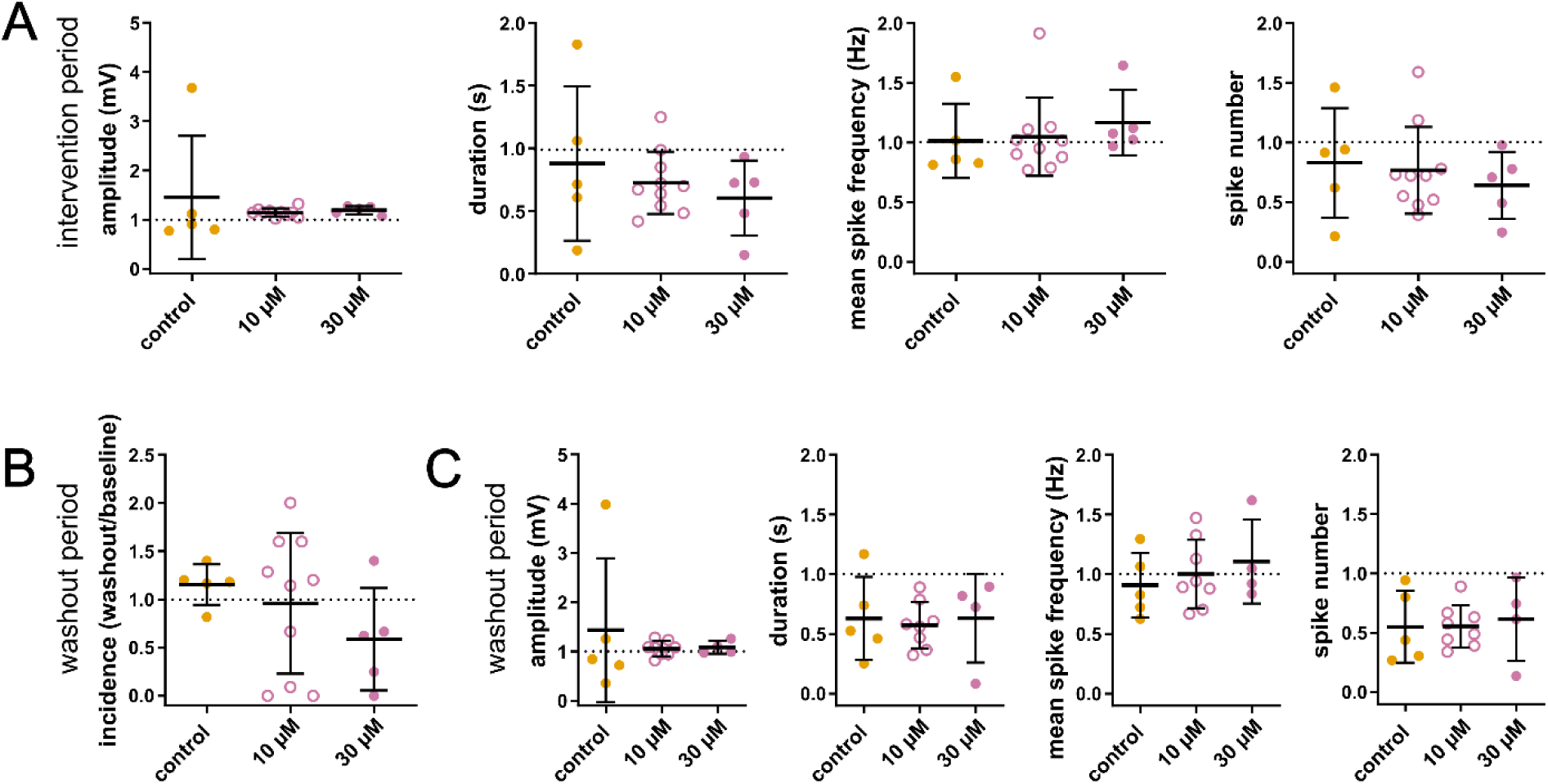
Effects of different concentrations of LY3130481 on seizure-like events *in vitro* in mice induced by 4-AP. Scatterplots show mean values ± standard deviation. All values are normalized to baseline values, meaning intervention/baseline for intervention period and washout/baseline for washout period. Statistics were done by Kruskal-Wallis test and Dunn’s multiple comparisons test. (A) Scatterplots of SLE amplitude, duration, mean spike frequency and spike number during intervention period compared between control group, LY3130481 10 µM and LY3130481 30 µM. (B) Scatterplots of SLE incidence during washout period. (C) Scatterplots of SLE amplitude, duration, mean spike frequency and spike number during washout period.

